# Age-dependent chemotherapy response and vitamin D impact in paired patient-derived normal and tumor colorectal organoids

**DOI:** 10.64898/2025.12.15.694334

**Authors:** Asunción Fernández-Barral, Antonio Barbáchano, Silvia Rodríguez-Marrero, Luis del Peso, Alfonso Herreros-Cabello, Nuria Rodríguez-Salas, Isabel Prieto, Aurora Burgos, Miguel León-Arellano, Damián García-Olmo, Tomás Olleros, José Manuel González-Sancho, María Jesús Larriba, Alberto Muñoz

**Author notes:** Indicates equal contribution (Asunción Fernández-Barral, Antonio Barbáchano). Correspondence: Asunción Fernández-Barral or María Jesús Larriba, Instituto de Investigaciones Biomédicas Sols-Morreale, Arturo Duperier, 4. 28029 Madrid. Spain /.

## Abstract

Early-onset colorectal cancer (EO-CRC, <50 years) incidence is rising worldwide. Using paired patient-derived colorectal normal and tumor organoids, we demonstrate the therapeutic window of three standard chemotherapies. Normal organoids were more resistant than tumor organoids to 5-fluorouracil (5-FU) and oxaliplatin in both EO-CRC and late-onset CRC (LO-CRC) patients. However, for SN38, this therapeutic window was observed in LO-CRC but not in EO-CRC patients, revealing age-dependent differences in drug resistance. We also evaluated the effect of calcitriol, the active vitamin D metabolite with anti-CRC activity, on drug sensitivity. Calcitriol reduced the cytotoxicity of 5-FU and SN38 in normal organoids regardless of patient age, while it selectively decreased their cytotoxicity in EO-CRC but not in LO-CRC tumor organoids. This protective effect correlated with calcitriol antiproliferative action and transcriptional effects on drug-metabolism pathways. These findings identify clinically relevant age-dependent differences in chemotherapy response and support vitamin D-based strategies to design age-tailored precision treatments.

## INTRODUCTION

Colorectal cancer (CRC) is a leading neoplasia in terms of incidence and mortality. Paradoxically, the incidence of CRC in patients less than 50 years of age (early-onset CRC, EO-CRC) is increasing for not well understood causes, while incidence in patients aged ≥50 years (late-onset CRC, LO-CRC) is declining, probably due to implementation of screening programs targeting preferentially aged populations and improved treatments ^1–4^. EO-CRC is usually diagnosed at advanced stages and is characterized by distinctive genetic, molecular and microbiome profiles, as well as more aggressive clinical behavior that may be due to delay in diagnosis. However, there are no age-specific treatment protocols for CRC patients ^2, 5–10^.

Organoids are considered an advantageous preclinical model for cancer modeling and precision therapy ^11–14^, in particular to assay the activity of antitumor drugs, with respect to classically used immortal cell lines and animal models ^15^. Aiming at a personalized anticancer therapy, organoids may provide functional data to enrich the genomic information obtained from mutational tumor analysis. As for CRC, organoids are being used in drug discovery and toxicity studies, and for the personalized treatment of CRC patients, although methodology needs to be improved ^16–20^. Of note, tumor organoids derived from EO-CRC patients have revealed a striking diversity of molecular phenotypes and heterogeneous genetic background ^21^.

A large number of epidemiological studies indicate that vitamin D deficiency is a risk factor for developing and dying from CRC ^22, 23^, and many supplementation studies and clinical trials support a protective action of vitamin D against CRC ^24–27^. Interestingly, high total vitamin D intake is associated with decreased risk of EO-CRC in a cohort of young women ^28^. Concordantly, the active vitamin D metabolite 1α,25-dihydroxyvitamin D_3_ (calcitriol) displays a battery of anti-CRC mechanisms in experimental systems ^29–33^. Among these mechanisms, calcitriol has been proposed to potentiate cancer therapy and to reverse drug-resistance in bladder, gastric, breast, glioblastoma, liver, ovarian, pancreatic and other cancer cell types ^34–38^. Furthermore, calcitriol promotes cell sensitivity to 5-fluorouracil (5-FU) in human colon carcinoma cells by downregulating thymidylate synthase (5-FU molecular target), and the anti-apoptotic protein survivin^39^.

Importantly, most reported studies on the action of calcitriol on CRC were performed using immortal long-term cultured carcinoma cell lines, which do not properly reproduce the *in vivo* situation. Tumors are thought to originate from genetically and epigenetically altered tissue resident stem cells (known as cancer stem cells, CSC) which are also proposed as responsible for chemotherapy resistance and tumor relapse ^40^. Unlike 2D monolayers, 3D patient-derived organoids (PDOs) better recapitulate tissue architecture and function. Therefore, to gain more relevant information we have now studied the response to three anticancer drugs currently used in the clinic against CRC in patient-derived tumor organoids generated from CSC. Moreover, we used paired organoids from normal tissue from the same CRC patient to examine the therapeutic window of drug activity. This strategy allows the evaluation of both the efficacy and toxicity of anticancer treatments within the same genetic background, something that is not possible with other preclinical models. In addition, to investigate putative different age-dependent drug sensitivities we have separately analyzed the response to drugs in organoids from EO-CRC and LO-CRC patients. Lastly, we examined the effect of calcitriol on drug activity in normal and tumor PDOs across age groups.

## RESULTS

### Colorectal tumor PDOs are more sensitive to chemotherapy than normal PDOs

To analyze the chemosensitivity of colorectal PDOs, we established paired normal and tumor organoids from 10 CRC patients equally distributed by age (5 EO-CRC and 5 LO-CRC; Supplementary Table 1 and Fig. 1a and performed a low-throughput drug assay as described in Methods. Both normal and tumor PDOs were treated for four days with three clinically relevant chemotherapeutic agents for CRC: 5-FU, oxaliplatin and SN38 (Fig. 1b). Normal PDOs exhibited a variable response to the three drugs (Supplementary Fig. 1a). This heterogeneity is reflected in the heatmap of normalized *z*-scores of AUC (Supplementary Fig. 1b). Dose-response curves of mean values for each drug revealed SN38 as the most cytotoxic, whereas 5-FU and oxaliplatin exerted a comparable lower cytotoxicity (Supplementary Fig. 1c). Similar results were obtained with tumor PDOs, which also displayed variable responses to chemotherapy (Supplementary Fig. 1d and e), with the highest sensitivity to SN38 (Supplementary Fig. 1f). Since the cytotoxic activity of chemotherapeutic agents often relates to cell proliferation, we studied whether PDO growth rates were linked to their drug sensitivity. Notably, no significant correlations were found between cell proliferation rates of normal and tumor PDOs and their AUC values that could explain the observed differential drug chemosensitivities among patients (Supplementary Fig. 2).

**Fig. 1:**
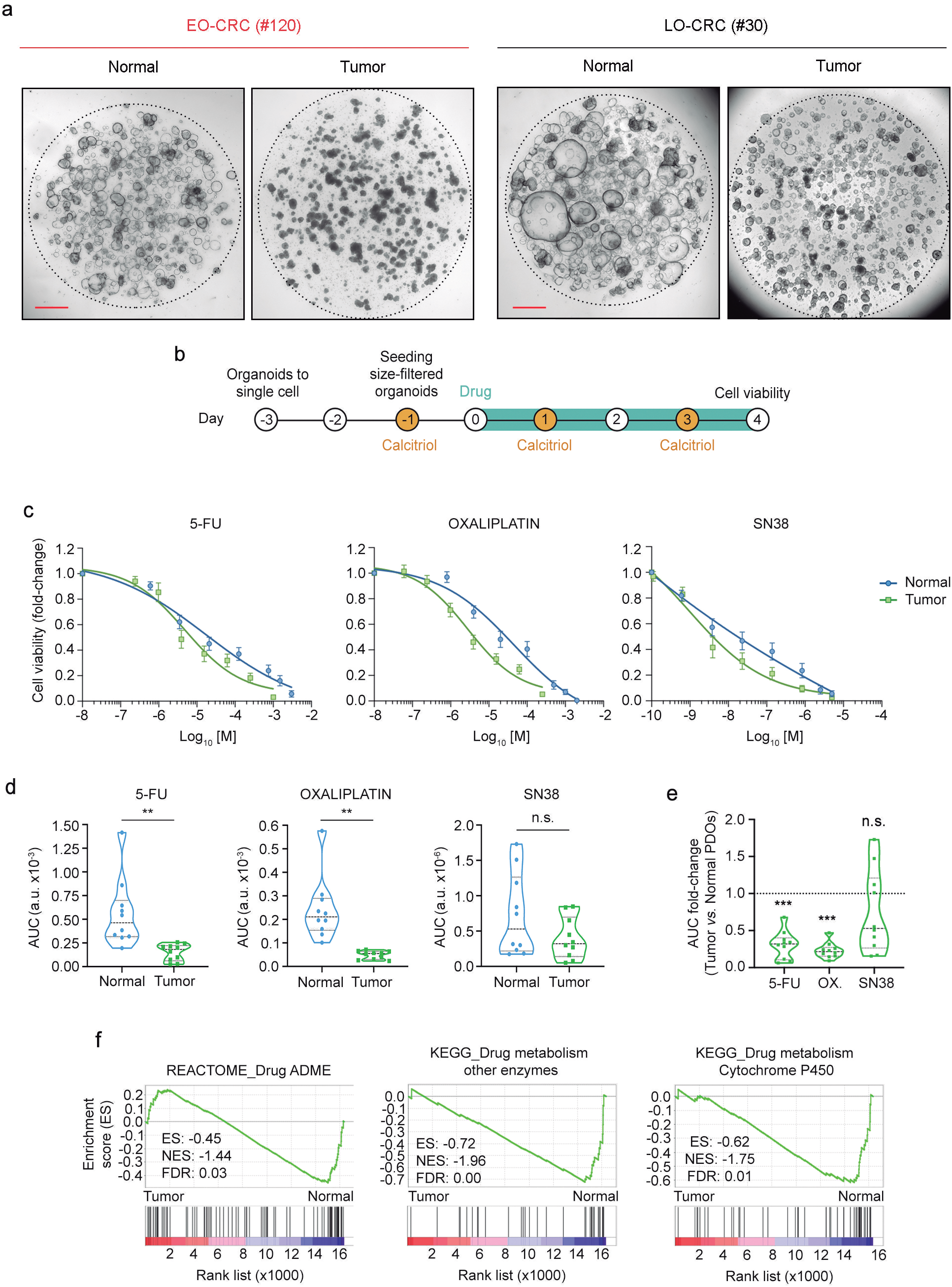
Colorectal normal and tumor PDOs display different drug sensitivity. **a** Representative bright-field microscopy images of paired normal and tumor organoids from EO-CRC (#120) and LO-CRC (#30) patients. Images correspond to organoid cultures during routine maintenance, prior to drug assays. Scale bar, 500 µm. **b** Scheme of the drug assay protocol. **c** Dose-response curves of paired normal and tumor PDOs treated with 5-FU, oxaliplatin or SN38. Mean values+/- SEM are plotted (n = 10). **d** Violin graphs showing the AUC values of drug assays from Supplementary Fig. 1a and d. Statistical analysis was performed by paired-test (n.s., non-significant; **P<0.01). **e** Violin graph depiciting the AUC fold-change values of tumor PDOs relative to their paired normal PDOs using data from (d). Discontinuous line represents normal PDOs. Statistical analysis was carried out using a one sample t-test (n.s., non-significant; ***P<0.001). **f** GSEA showing the positive association between drug metabolism-related gene signatures and the transcriptomic profile of normal organoids from CRC patients identified in a previous study^41^.

Comparison of dose-response curves of mean values from both types of PDOs revealed that tumor PDOs are significantly more sensitive to 5-FU and oxaliplatin than normal PDOs (Fig. 1c and d), thereby demonstrating the utility of human organoids to identify the therapeutic window *in vitro*. A similar trend was observed for SN38, that did not reach statistical significance probably due to the variable response of normal PDOs to this drug (Fig. 1c and d). This result was confirmed when analyzing the AUC fold-change between paired normal and tumor cultures, a comparison only possible using paired PDOs (Fig. 1e). Interestingly, GSEA of previously published RNA-seq data from paired colon normal and tumor organoids from CRC patients ^41^ showed that the gene expression profile of normal PDOs was positively associated with drug metabolism-related signatures (Fig. 1f). These data constitute a potential explanation for the reduced sensitivity of normal PDOs to chemotherapeutic drugs.

### Age-dependent differences in PDOs chemosensitivity

Considering the increasing clinical relevance of EO-CRC, we next evaluated PDO drug responses according to patient age. First, we found that normal organoids from EO-CRC and LO-CRC patients were less sensitive to 5-FU and oxaliplatin than tumor organoids, supporting the existance of a therapeutic window *in vitro* for these two drugs across age (Fig. 2a and b). Second, age-based stratification showed that EO-CRC and LO-CRC PDOs displayed a different response to SN38: in young patients, both normal and tumor PDOs exhibited similar sensitivity to SN38, whereas in aged patients a trend toward higher sensitivity of tumor PDOs was observed (Fig. 2b). This difference became significant when analyzing the AUC fold-change between paired normal and tumor PDOs, indicating that the therapeutic window for SN38 is maintained in LO-CRC PDOs but it is lost in organoids from EO-CRC patients (Fig. 2c).

**Fig. 2:**
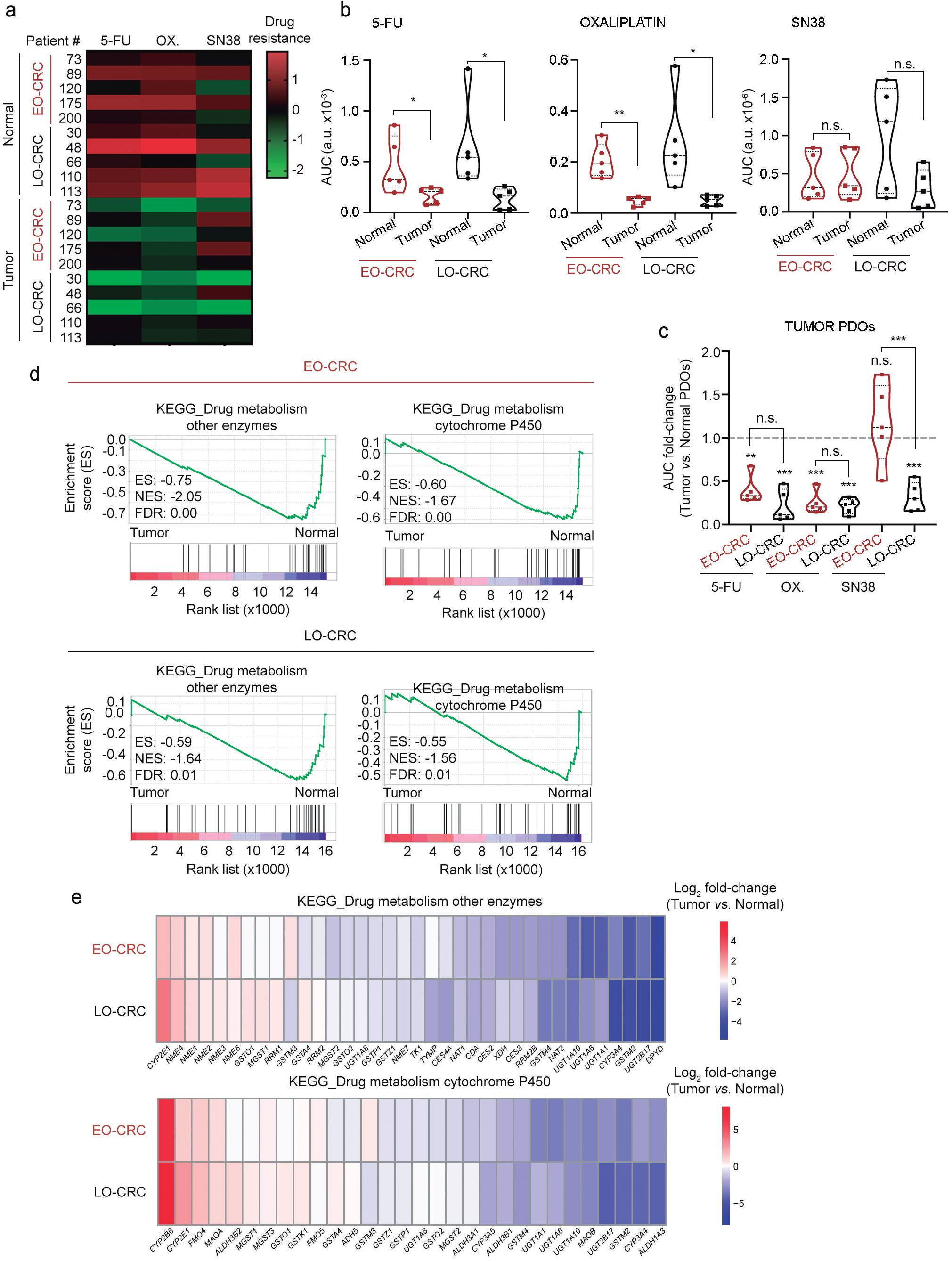
Age-dependent drug sensitivity of colorectal normal and tumor PDOs. **a** Heatmap illustrating the z-score of drug resistance of age-grouped normal and tumor PDOs treated with 5-FU, oxaliplatin or SN38. **b** Violin graphs showing the AUC values of age-grouped normal and tumor PDOs treated with 5-FU, oxaliplatin or SN38 from Supplementary Fig. 1a and d. Statistical analysis was performed by paired t-test (n.s., non-significant; *P<0.05; **P<0.01). **c** Violin graph representing the AUC fold-change values of tumor organoids from EO-CRC and LO-CRC patients relative to their paired normal PDOs using data from (b). Discontinuous line corresponds to normal PDOs. Differences between tumor and paired normal PDOs were evaluated by one-sample t-test, and comparisons of therapeutic windows between age groups were performed using one-way ANOVA with Sidak’s post-test (n.s., non-significant; **P<0.01; ***P<0.001). **d** GSEA showing the positive association between the transcriptomic profile of normal organoids from EO-CRC and LO-CRC patients and gene signatures related with drug metabolism. **e** Heatmaps illustrating the differential gene expression (tumor vs. normal PDOs) in EO-CRC and LO-CRC patients for the genes included in the drug metabolism signatures identified in (d).

In addition, we performed an RNA-seq analysis of paired normal and tumor PDOs from 5 EO-CRC and 6 LO-CRC patients (Supplementary Table 1). This study revealed that normal organoids from both young and aged patients exhibit expression profiles enriched in drug metabolism-related signatures (Fig. 2d and e), which is in line with the results from our previously published RNA-seq analysis of PDOs lacking age stratification (Fig. 1f). No age-related differences were observed in the enrichment of these signatures (Fig. 2d and e).

### Calcitriol modulates drug resistance of normal and tumor PDOs

Next, we sought to investigate the effect of calcitriol on the drug sensitivity of normal and tumor PDOs. First, we examined the responsiveness to calcitriol of the two types of organoids. As shown in Supplementary Fig. 3a, both normal and tumor PDOs, regardless of patient age, expressed comparable levels of *VDR* RNA encoding the vitamin D receptor. Accordingly, *CYP24A1*, the most responsive calcitriol target gene, which encodes the enzyme responsible for its degradation, was upregulated by calcitriol in normal and tumor PDOs from both age groups (Supplementary Fig. 3b). Concordantly with reports of *CYP24A1* overexpression in human colorectal tumors ^42, 43^, the basal expression of *CYP24A1* was higher in tumor PDOs than in normal PDOs (Supplementary Fig. 3b).

Paired normal and tumor PDOs were treated with 5-FU, oxaliplatin or SN38 in the presence or absence of calcitriol as described in Fig. 1b. Strikingly, in normal PDOs calcitriol decreased the sensitivity to the three drugs (Fig. 3a-c), suggesting a protective effect on healthy colon mucosa upon chemotherapy. Likewise, in tumor PDOs the sensitivity to oxaliplatin and SN38, but not to 5-FU, was also reduced by calcitriol (Fig. 3d-f). In line with this, GSEA of RNA-seq data including calcitriol-treated PDOs revealed that drug metabolism-related signatures were significantly enriched in both normal and tumor PDOs upon calcitriol treatment regardless of age (Fig. 4a).

**Fig. 3:**
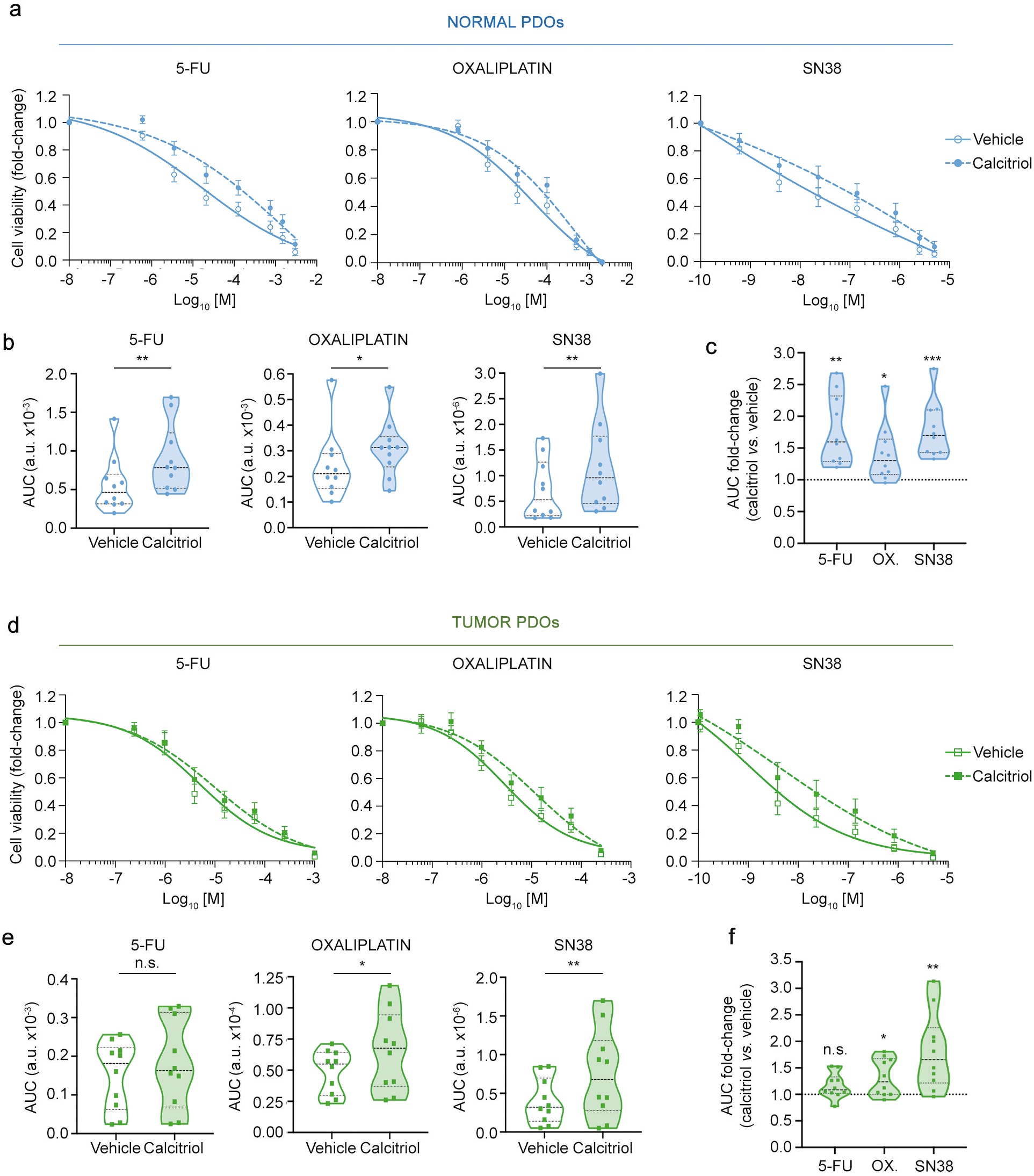
Calcitriol modulates drug cytoxicity in colorectal normal and tumor PDOs. **a** Dose-response curves of normal PDOs to 5-FU, oxaliplatin and SN38 in the presence of 100 nM calcitriol or vehicle (ethanol) for 96 h. Mean values+/- SEM are plotted (n = 10). **b** Violin graphs showing the AUC values of drug assays from (a). **c** Violin graph displaying the AUC fold-change values of calcitriol-vs. vehicle-treated normal PDOs using data from (b). Discontinuous line represents vehicle-treated normal PDOs. **d** Dose-response curves of tumor PDOs to 5-FU, oxaliplatin and SN38 in the presence of 100 nM calcitriol or vehicle (ethanol) for 96 h. Mean values+/- SEM are plotted (n = 10). **e** Violin graphs showing the AUC values of drug assays from (d). **f** Violin graph displaying the AUC fold-change values of calcitriol-*vs.* vehicle-treated tumor PDOs using data from (e). Discontinuous line represents vehicle-treated tumor PDOs. Statistical analyses of (b) and (e) were performed by paired I-test (n.s., non-significant; *P<0.05; **P<0.01) and of (c) and (f) by one-sample I-test (n.s., non-significant; *P<0.05; **P<0.01; ***P<0.001).

**Fig. 4:**
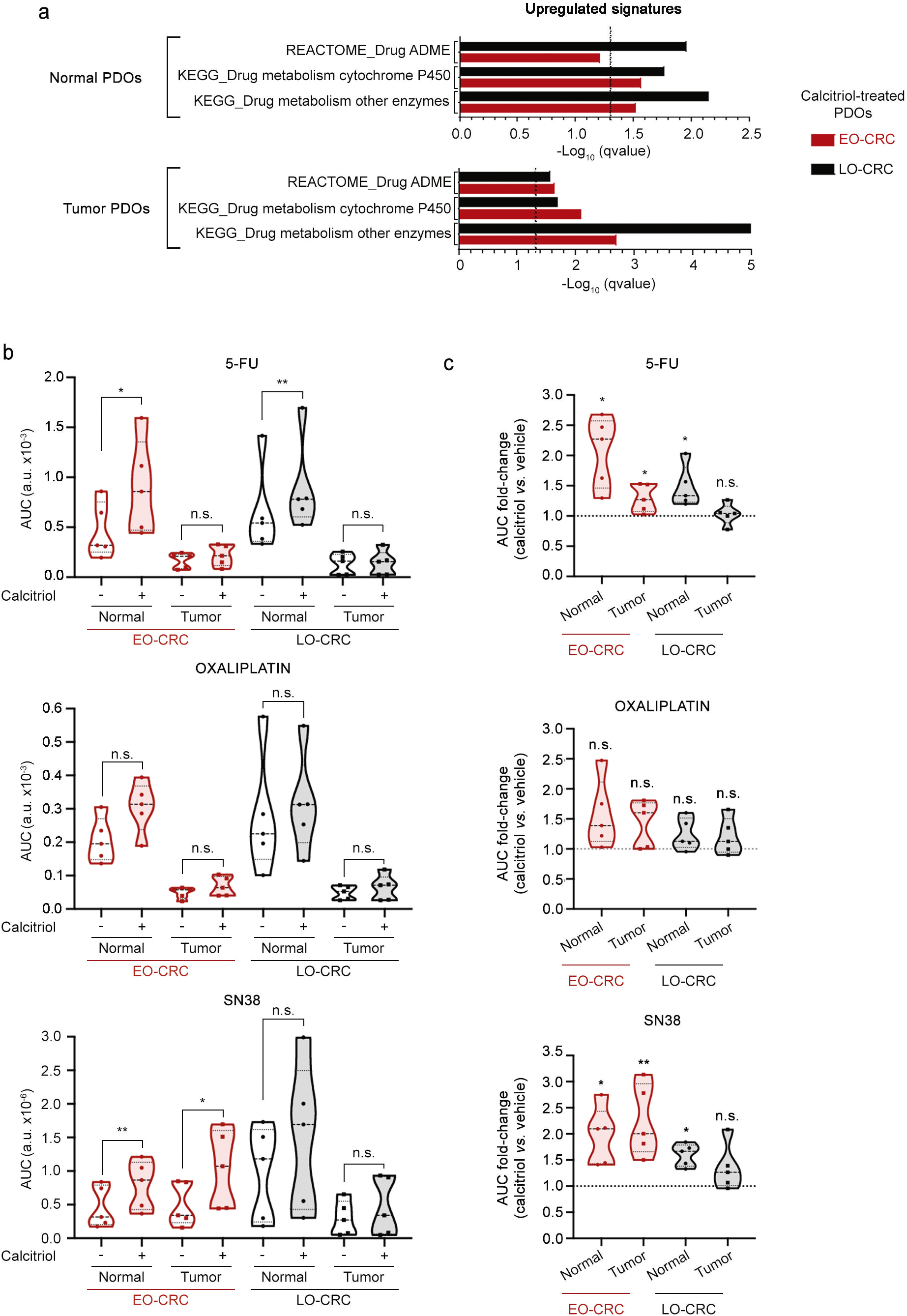
Differential effect of calcitriol on drug cytotoxicity in EO-CRC and LO-CRC tumor PDOs. **a** Graph showing the association identified by GSEA of RNA-seq data between drug metabolism-related signatures and the gene expression profiles of calcitriol-treated normal and tumor organoids from EO-CRC and LO-CRC patients. Discontinous line represents the cut-off for statistical significance. **b** Violin graphs showing the AUC values of age-grouped normal and tumor PDOs treated with 5-FU, oxaliplatin or SN38 in the presence of calcitriol (100 nM) and vehicle (ethanol) for 96 **h**. Statistical analysis was performed by paired I-test (n.s., non-significant; *P<0.05; **P<0.01). **c** Violin graphs re-presenting the AUC fold-change values of calcitriol-vs. vehicle-treated normal and tumor organoids from EO-CRC and LO-CRC patients using data from (b). Discontinuous line corresponds to vehicle-treated PDOs (n.s., non-significant; *P<0.05; **P<0.01).

Dose-response analyses of age-stratified normal and tumor PDOs showed that patient age defines the response to chemotherapy under calcitriol treatment. Calcitriol diminished the cytotoxic activity of 5-FU and SN38 in normal organoids from both EO-CRC and LO-CRC patients (Supplementary Fig. 4, Fig. 4b and c). Strikingly, calcitriol had age-dependent differential effects on the sensitivity of tumor PDOs to 5-FU and SN38: it diminished their cytotoxicity in young patients but did not alter drug sensitivity in organoids derived from aged patients (Supplementary Fig. 4, Fig. 4b and c. These age-related effects were further supported by the analysis of AUC fold-changes between calcitriol- and vehicle-treated PDOs, where the reduction exerted by calcitriol on the sensitivity of LO-CRC normal PDOs to SN38 and that of EO-CRC tumor PDOs to 5-FU reached statistical significance (Fig. 4c). Together, these results suggest that calcitriol cotreatment may favor a wider therapeutic window for 5-FU and SN38 specifically in LO-CRC patients. In contrast, calcitriol did not modify oxaliplatin sensitivity in normal or tumor PDOs across age groups (Supplementary Fig. 4, Fig. 4b and c), even though this effect was observed when age stratification was not considered (Fig. 3b, c, e, f).

### Age-dependent differential antiproliferative effect of calcitriol in tumor PDOs

We next explored potential age-dependent differences in cell proliferation inhibition by calcitriol in paired normal and tumor PDOs. Calcitriol consistently inhibited cell proliferation in normal PDOs, while, in contrast, it showed substantial interindividual variability in tumor PDOs (Fig. 5a). Stratification by age revealed that calcitriol reduced cell proliferation in EO-CRC and LO-CRC normal PDOs and also in EO-CRC tumor PDOs, but had no effect in LO-CRC tumor PDOs (Fig. 5b). In line with these results, GSEA of RNA-seq data revealed a significant negative association between the gene expression profile of calcitriol-treated EO-CRC tumor PDOs and cell proliferation-related signatures, an association that was not observed in LO-CRC tumor PDOs (Fig. 5c). These findings indicate that the calcitriol-induced proliferative arrest in tumor PDOs is restricted to young patients. Of note, while no significant correlations between cell proliferation and drug response were observed in vehicle-treated PDOs (Supplementary Fig. 2), significant positive correlations were detected between the antiproliferative effect of calcitriol and the increase in drug resistance exerted by calcitriol in both normal and tumor (except for oxaliplatin) PDOs (Fig. 5d).

**Fig. 5:**
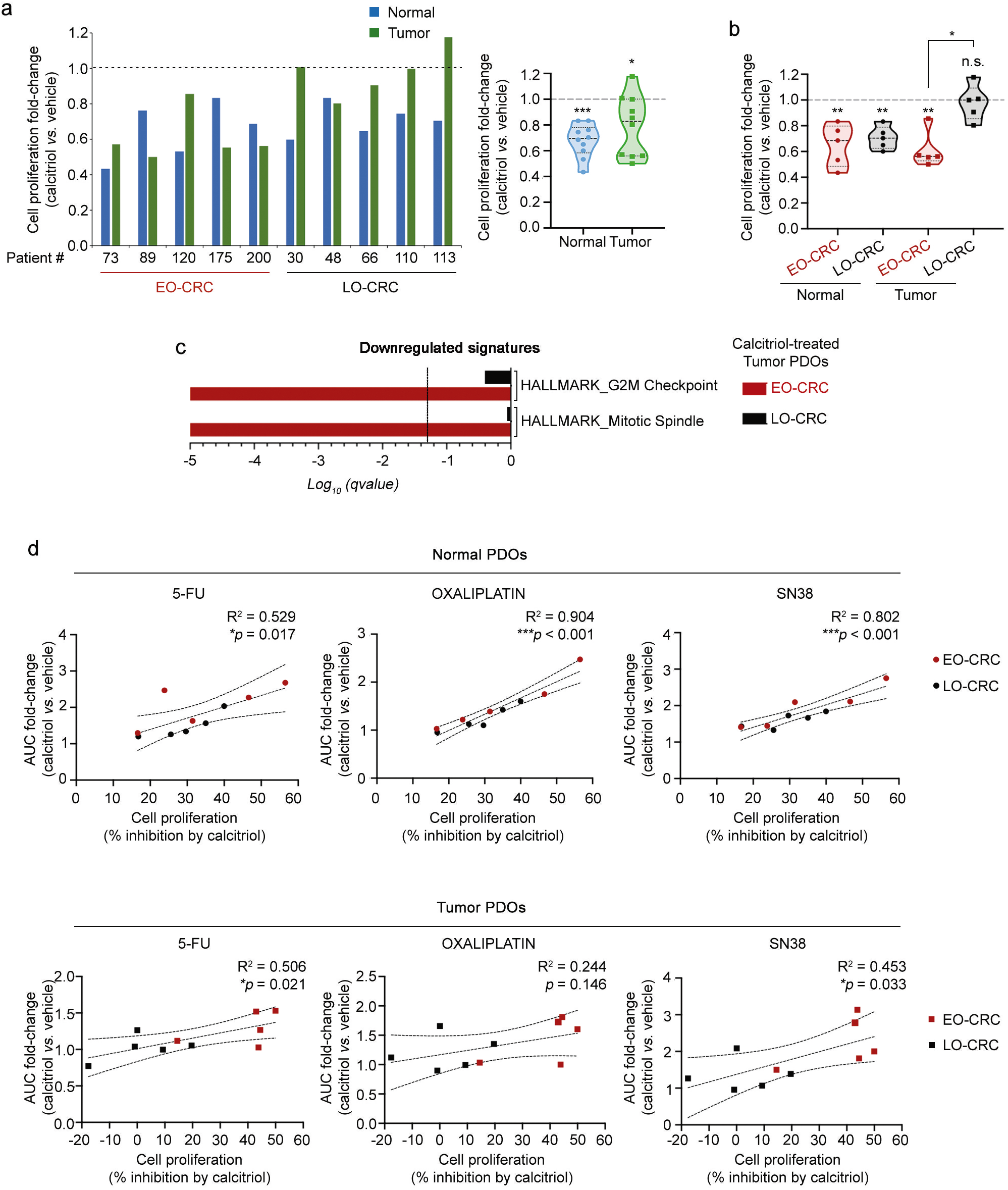
Relationship between drug cytotoxicity and cell proliferation in colorectal normal and tumor PDOs treated with calcitriol. **a** Cell proliferation assays of paired normal and tumor PDOs treated with 100 nM calcitriol or vehicle (ethanol) for 96 h. Bar (left panel) and violin (right panel) graphs showing the effect of calcitriol. Statistical analysis was performed by one sample I-test (*P<0.05; ***P<0.001). b Violin graph illustrating the calcitriol-induced fold-change in cell proliferation in PDOs from (a) stratified by age. Differences relative to vehicle-treated organoids were evaluated by one-sample-test (n.s., non-significant; **P<0.01), and comparison between EO-CRC and LO-CRC tumor PDOs was performed using the upaired-test (*P<0.05). Discontinuous lines in (a) and (b) correspond to vehicle-treated organoids. c Graph showing the negative association identified by GSEA of RNA-seq data between cell proliferation-related signatures and the gene expression profiles of calcitriol-treated tumor organoids from EO-CRC and LO-CRC patients. Discontinous line represents the cut-off for statistical significance. d Scattergrams displaying the correlations between the effect of calcitriol on cell proliferation and its impact on the cytotoxicity of 5-FU, oxaliplatin and SN38 in normal (upper panels) and tumor (lower panels) organoids from EO-CRC and LO-CRC patients. Statistical correlation was evaluated with Pearson’s test (*P<0.05; ***P<0.001).

In summary, our results show that calcitriol exerts distinct, age-dependent effects on the sensitivity of CRC PDOs to 5-FU and SN38 antitumoral drugs. In young EO-CRC patients, calcitriol increases drug resistance in both normal and tumor PDOs, thereby preserving the therapeutic window observed in its absence. Contrarily, in aged LO-CRC patients, calcitriol selectively increases drug resistance in normal PDOs but not in tumor PDOs, resulting in a widened therapeutic window.

## DISCUSSION

To our knowledge, this is the first study comparing the response to antitumor drugs of paired normal and tumor colorectal organoids from young and aged CRC patients. Our results show that normal organoids derived from EO-CRC and LO-CRC patients are less sensitive to currently used drugs such as 5-FU and oxaliplatin than their paired tumor counterparts, which define the existence of a therapeutic window *in vitro* that may be the basis of their clinical utility. We show that this reduced sensitivity is likely attributable to the elevated expression of drug metabolism-related genes in normal PDOs, rather than an age-dependent trait. In the case of SN38, the lower sensitivity of normal *vs.* tumor organoids only applies to LO-CRC organoids, suggesting higher intestinal toxicity of this drug in young than in aged patients. In addition, we report that tumor organoids from EO-CRC and LO-CRC patients display similar sensitivity to each of the three antitumor drugs commonly used in clinical practice. This is in line with data regarding response to chemotherapy of these two groups of patients ^2, 7, 44^.

Organoids derived from both normal and tumor tissues of the same patient constitute a unique experimental platform, as they enable the evaluation of drug efficacy and toxicity within a genetically paired system. This dual organoid approach is particularly powerful because it allows the assessment of antitumor drug selectivity, distinguishing their cytotoxic effects on malignant cells from those on healthy epithelia, which cannot be achieved with conventional cancer cell lines. Therefore, the present study not only provides insights into the age-dependent drug response of CRC tumors, but also shows the potential of PDOs to model therapeutic selectivity in a personalized manner. Additionally, testing drug sensitivity in both types of PDOs allows to identify which agents are more effective against tumor cells and also to estimate doses that could minimize adverse effects on healthy (intestinal) tissue. Thus, this *in vitro* therapeutic window could inform personalized treatment strategies and guide dose optimization in the clinical setting. Although translation from *in vitro* to *in vivo* remains challenging, these assays provide a useful framework to explore such strategies.

We performed drug assays on fully formed organoids to preserves the 3D structure and cellular interactions. This strategy better recapitulates the *in vivo* tissue architecture and provides a more realistic assessment of drug efficacy. In addition, the determination of optimal organoid size and density helps to minimize variability and ensures reproducibility across experiments. This methodological refinement enhances assay robustness and distinguishes the drug assay performed from those developed with traditional 2D cultures or with other 3D systems.

Interestingly, our results reveal that the active vitamin D metabolite calcitriol diminishes the cytotoxicity of 5-FU and SN38 in colorectal normal organoids from both young and aged patients. This result suggests a possible reduction in intestinal side effects/toxicity by these two drugs under an appropriate vitamin D status. Calcitriol reduces also the cytotoxic action of 5-FU and SN38 in EO-CRC but not in LO-CRC tumor organoids, which points to a preferential beneficial effect of the combined treatment of these two drugs and vitamin D compounds in aged CRC patients. The capacity of calcitriol to diminish drug cytotoxicity in colorectal stem cell-derived organoids contradicts data obtained in established carcinoma cell lines indicating the potentiation by vitamin D compounds of the activity of antitumor drugs ^34–38^. Possible explanations for this discrepancy include the usual highly malignant features of cell lines that are established with low success from human tumors and the accumulation of karyotypic aberrations during their long-term culture. Supporting this idea, calcitriol has distinct effects on gene expression in colon carcinoma cells and colon tumor organoids, as it is exemplified by the repression of Wnt/β-catenin target genes in the former but not in latter system ^41, 45^.

Calcitriol has been shown to display a wide range of protective actions against CRC that include the inhibition of cancer cell proliferation and invasion, angiogenesis and metastasis, the promotion of cell differentiation and apoptosis, and the potentiation of immune responses ^24, 32, 33^. Accordingly, VDR expression is a good prognostic biomarker and correlates with immune infiltration in CRC ^46, 47^. Moreover, recent data show that vitamin D positively modulates intestinal microbiome composition and augments immune-dependent resistance to cancers in mice, while in humans the expression of calcitriol-induced genes correlates with improved responses to immune checkpoint inhibitors ^48^. In line with all these data, the majority of recent reviews and meta-analyses of epidemiological and supplementation studies support a beneficial effect of an adequate vitamin D status in CRC ^24–27^. Here, we report that calcitriol reduces cell proliferation in colorectal PDOs, an effect that is associated with decreased cytotoxicity of antitumor drugs. Of note, this action takes place in normal organoids from all-age patients and in tumor PDOs from EO-CRC patients, but only in a subset of LO-CRC tumor PDOs. These data are compatible with the reported global anticancer action of calcitriol and, moreover, suggest that the combination of anticancer therapies with an adequate vitamin D status or with vitamin D compounds may enhance the therapeutic window of the antitumor drugs preferentially in aged patients.

A limitation of our study is the pure epithelial nature of the organoids used, as several features of cancer cells, including their response to drugs, are modulated by the surrounding stroma ^49^. Thus, cancer-associated fibroblasts have been shown to induce therapy resistance in cancer cells and CRC organoids ^50, 51^. Moreover, the organoid system has other intrinsic and experimental limitations including moderate success rate and difficult standardization, which still need to be optimized ^52, 53^. Nevertheless, interpatient variability in drug response across PDOs closely mirrors clinical diversity, representing one of the major strengths of the organoid system for precision medicine applications. Moreover, PDOs retain a significant degree of intratumoral heterogeneity, preserving diverse cellular populations and subclonal architectures that are present in the original tumor. In addition, organoid apical-basal orientation might limit certain applications, such as co-culture with microorganisms or nutrient absortion assays, due to the difficulty in accessing to the apical pole ^54^. However, this feature is less critical in tumor PDOs, which often display compact and non-polarized structures, making drug assays largely unaffected. Overall, PDOs are considered a valuable drug screening platform and a promising system to predict patient response to treatment in a variety of cancer types ^17, 55^. Consistent with this, a good correlation between PDOs chemosensitivity and CRC patient response has been reported to some, but not all, chemotherapy drugs ^20, 56, 57^, and recent studies show that PDOs sensitivity correlates with the response of metastatic CRC patients to 5-FU, oxaliplatin and irinotecan^58^.

In conclusion, this study indicates that colorectal normal and tumor PDOs exhibit age-dependent sensitivity to antitumor drugs commonly used for CRC and to the modulatory action of calcitriol on drug activity. Our results also support an age-independent role for vitamin D in attenuating the intestinal side effects of chemotherapy and highlight the potential value of designing combinations of chemotherapy and vitamin D compounds according to patient age.

## METHODS

### Human samples

The collection of paired normal and tumor CRC patient tissue samples for the establishment of organoids was approved by the Ethics Committee of Hospital Universitario La Paz (HULP-PI-1425, HULP-PI-3196, HULP-PI-4462). Fresh human tissues biopsies were obtained from CRC patients undergoing surgery or diagnostic colonoscopy between 2014 and 2023 and provided by IdiPAZ Biobank (PT23/00028), integrated into the Spanish Biobank Network (https://www.isciiibiobanksbiomodels.es/). All patients provided written informed consent.

### Generation and culture of patient-derived organoids

Colorectal normal and tumor PDOs were generated as previously described ^41^. Briefly, human colon normal biopsies, collected from areas distant to the tumor, were incubated in rotation with an antibiotic mixture (Primocin [Invivogen, CA, USA], gentamicin and fungizone [Thermo Fisher Scientific, MA, USA]) for 1 h at room temperature (RT). Biopsies were then cut into small fragments and incubated under slow rotation twice for 5 min with 10 mM dithiothreitol (DTT [Tocris, Bristol, UK]) at RT followed by incubation with 8 mM EDTA for 5 min at RT and 60 min at 4°C. After several PBS washes to remove EDTA, samples were gently shaken in fresh PBS to release isolated crypts. Pelleted crypts were washed in washing buffer (Advanced DMEM/F12, 10 mM HEPES, and 10 mM Glutamax [Thermo Fisher Scientific]), embedded in Matrigel (Corning, NY, USA) and seeded as drops onto pre-warmed 6-well plates. Following Matrigel polymerization, normal culture medium (Supplementary Table 2) was added. To establish patient-derived tumor organoids, tumor tissues were incubated with the same antibiotic mixture as normal samples. Biopsies were enzymatically digested with 1 mg/mL collagenase type IV (Sigma-Aldrich, MA, USA) in PBS for 30 min under rotation at 37°C. Cell suspension was passed through an 18 G syringe and subsequently filtered through a 100 μm mesh filter. Erythrocytes were lysed by incubating the cell suspension with 157 mM NH_4_Cl for 5 min. Single cells were then embedded in Matrigel and plated onto pre-warmed 12-well dishes. Following Matrigel polymerization, tumor culture medium (normal medium lacking L-WNR conditioned medium, RSPO1, nicotinamide, and supplemented with 10 μM Y27632 [Tocris]) (Supplementary Table 2) was added. Culture medium for both normal and tumor organoids was refreshed every other day.

PDOs were passaged according to a previously described protocol ^59^. Briefly, organoids were incubated for 30 min with 1 mg/mL dispase (Thermo Fisher Scientific) at 37°C. Matrigel-embedded organoids were collected with a scraper and centrifuged at 269 x *g* for 5 min. Pelleted organoids were resuspended in a 1:4 TrypLE Express (Gibco, NY, USA) in PBS solution for 5 min at 37°C, followed by mechanical disaggregation through a 21 G syringe. After two washes in washing buffer, pelleted cells were embedded in Matrigel and seeded into culture dishes. Y27632 was maintained in both normal and tumor media until fragmented organoids or single cells generated complete organoids.

### Drug activity assays

Culture conditions for cell viability assays of normal PDOs were optimized from a protocol previously described by our group for tumor PDOs ^60^. Modifications were introduced to obtain a proportional relationship between luminescence signal and organoid number, through the adjustment of organoid size and seeding density. For this characterization prior to the assay, normal organoids devoid of Matrigel were sequentially filtered through 70 μm, 40 μm and 20 μm strainers (Pluriselect, Leipzig, Germany) to obtain two size-selected populations: 20-40 μm and 40-70 μm. Both fractions were resuspended in 1 mL culture medium, and organoid concentration was determined. Eight conditions (20-40 μm: 125, 250, 500, and 1000 organoids; 40-70 μm: 50, 100, 200, and 400 organoids) were tested in triplicate by seeding organoids into 96-well dishes containing 2% Matrigel. After four days of treatment, cell viability was measured as cellular ATP level using CellTiter-Glo 3D Luminescent Cell Viability Assay (Promega, WI, USA) following the manufacturer’s instructions. The optimal seeding density for normal PDOs was defined as 500 organoids of 20-40 μm per well, which ensures a proportional luminescence signal relative to the number of organoids seeded. Tumor PDOs were previously optimized (250 organoids of 20-40 μm per well) using an equivalent protocol ^60^.

Following patient characterization, full-grown paired normal and tumor PDOs were collected after removing Matrigel. Pelleted organoids were resuspended in washing buffer and filtered through 20 μm and 40 μm cell strainers. Size-selected organoids were centrifuged and resuspended in 500 μL of the corresponding culture medium. Organoid density was determined and adjusted to seed 250 tumor and 500 normal organoids per well, distributed in four 5 μL drops within a 24-well dish. For each drug, two complete 24-well dishes were seeded: one treated with calcitriol (100 nM) and the other with vehicle (ethanol). On the following day, organoids were treated in triplicate with seven drug concentrations plus a control (vehicle: water for oxaliplatin, and DMSO for SN38 and 5-FU). We used 5-FU, oxaliplatin and SN38 (active metabolite of irinotecan) (all from Sigma-Aldrich). Retreatment with calcitriol was performed every other day. After four days of drug exposure, cell viability was assessed using the CellTiter-Glo 3D Cell Viability Assay as previously described ^60^. Drug responses were quantified as the area under the curve (AUC) using GraphPad Prism v10 (GraphPad Software, CA, USA) by numerical integration (trapezoidal rule). For each drug, patient-specific AUC values were standardized to z-scores (*z = (LnAUC - μ) / σ*), where μ and σ denote the cohort mean and standard deviation, respectively. These *z*-scores, reflecting the relative drug resistance of each PDO, were visualized in a heatmap.

### Cell proliferation assays

Cell proliferation assays were carried out under similar conditions to the viability assays. Size-selected (20-40 μm) normal (500) and tumor (250) organoids were seeded in four 5 μL drops in 24-well dishes, in triplicate. After Matrigel polymerization, the corresponding culture medium was added and supplemented with either 100 nM calcitriol or vehicle (ethanol). Culture medium was replaced every other day together with calcitriol retreatment. After four days in culture, proliferation was determined using the CellTiter-Glo 3D Cell Viability Assay (Promega).

### RNA extraction and reverse transcription quantitative PCR (RT-qPCR)

Matrigel-embedded normal and tumor organoids were lysed by using TRIZOL (Thermo Fisher Scientific). Total RNA was purified using the NucleoSpin miRNA extraction kit (Machery-Nagel, Düren, Germany), and for cDNA retrotranscription the iScript cDNA Synthesis kit (Bio-Rad, CA, USA) was used. qPCR analyses were performed with the Taqman Universal PCR Master Mix (Applied Biosystems, CA, USA) using a FAM-labeled TaqMan probe for *CYP24A1* (Hs00167999_m1, Applied Biosystems). RNA expression values were normalized *vs.* the housekeeping gene *RPLP0* (H99999902_m1; VIC-labeled TaqMan probe; Applied Biosystems) using the comparative CT method. Additionally, we also used the Power SYBR Green PCR Master Mix (Applied Biosystems) and the following primers for *VDR* detection (forward 5′-AACGCTGTGTGGACATCGGC-3′; reverse 5′-GTCATGGCTTTCGTTGGACT-3′) and their values were normalized *vs*. the housekeeping gene *SDHA* (forward 5′-GGGAACAAGAGGGCATCTG-3′; reverse 5′-CCACCACTGCATCAAATTCATG-3′) using the comparative CT method. All qPCR analyses were performed in a CFX384 Touch Real-Time PCR Detection System (Bio-Rad). RNA from human SW480-ADH colon carcinoma cells were used to obtain a relative value to compare *VDR* and *CYP24A1* expression.

### RNA-sequencing (RNA-seq) assay

Paired normal and tumor organoids from 5 EO-CRC and 6 LO-CRC patients were seeded in 12-well culture dishes and 48 h later were treated with calcitriol (100 nM) or vehicle (ethanol) for 96 h. Medium and calcitriol treatment were refreshed every two days. Total RNA samples (1µg; RIN range 8.6-9.9) were converted into sequencing libraries with the NEBNext Ultra II Directional RNA Library Prep Kit for Illumina (New England Biolabs, Ipswich, MA, USA). Briefly, polyA+ fraction was purified and randomly fragmented, converted to double stranded cDNA and processed through subsequent enzymatic treatments of end-repair, dA-tailing, and ligation to adapters. Directional cDNA libraries, stranded in the antisense orientation were completed by PCR and pair-end sequenced on an Illumina NovaSeq X by following manufacturer’s protocols.

Raw FASTQ reads were aligned to the human GRCh38 v40 reference genome using HISAT2 (v2.1.0) with default parameters in paired-end mode. Alignment files (SAM) were converted to BAM and coordinate-sorted using SAMtools (v1.6). Gene-level quantification was performed with HTSeq-count (v0.11.3) using GRCh38 GTF annotations and stranded argument set to reverse. Genes with low expression were filtered using the filterByExpr function from the edgeR package (R v4.3.3). Differential expression analysis was conducted with the limma-voom pipeline, and p-values were adjusted for multiple testing using the Benjamini–Hochberg method.

Gene set enrichment analysis (GSEA) was carried out to evaluate the association between RNA-seq profiles and gene signatures enriched for proliferation and drug metabolism pathways obtained from MSigDB (www.gsea-msigdb.org). Heatmap analysis was done with R programming language (v4.5.0) using the package pheatmap.

### Statistical analyses

Statistical analyses were performed using GraphPad Prism v10. Significance was defined as \**P*<0.05, \*\**P*<0.01, and \*\*\**P*<0.001. The statistical test applied in each figure is indicated in its corresponding legend.

## DATA AVAILABILITY

RNA-seq data have been deposited in the European Nucleotide Archive (ENA) at EMBL-EBI under accession number PRJEB102090 (https://www.ebi.ac.uk/ena/browser/view/PRJEB102090). In addition, previously published transcriptomic data analyzed in this study are available at the Gene Expression Omnibus (GEO) database (accession number GSE100785) ^41^.

## ACKNOWLEDGEMENTS

We thank the patients and the Biobanks of IdiPAZ (PT23/00028) and Fundación Jiménez Díaz (PT23/00114), both integrated into the Spanish Biobank Network, for their collaboration.

## AUTHOR CONTRIBUTIONS

Conceptualization and design: AF-B, AB, MJL, AM. Methodology and data acquisition: AB, AF-B, SR-M. Bioinformatic analysis: LdP, AH-C. Resources: IP, ABu, NR-S, ML-A, DG-O, TO. Funding acquisition: NR-S, JMG-S, MJL, AM, TO. Writing – original draft: AM. Writing – review and editing: AF-B, AB, MJL, JMG-S. All authors approved and agreed to the final manuscript submission.

