## Supplementary Figure 1 for "Age-dependent chemotherapy response and vitamin D impact in paired patient-derived normal and tumor colorectal organoids"

Supplementary Fig. 1.

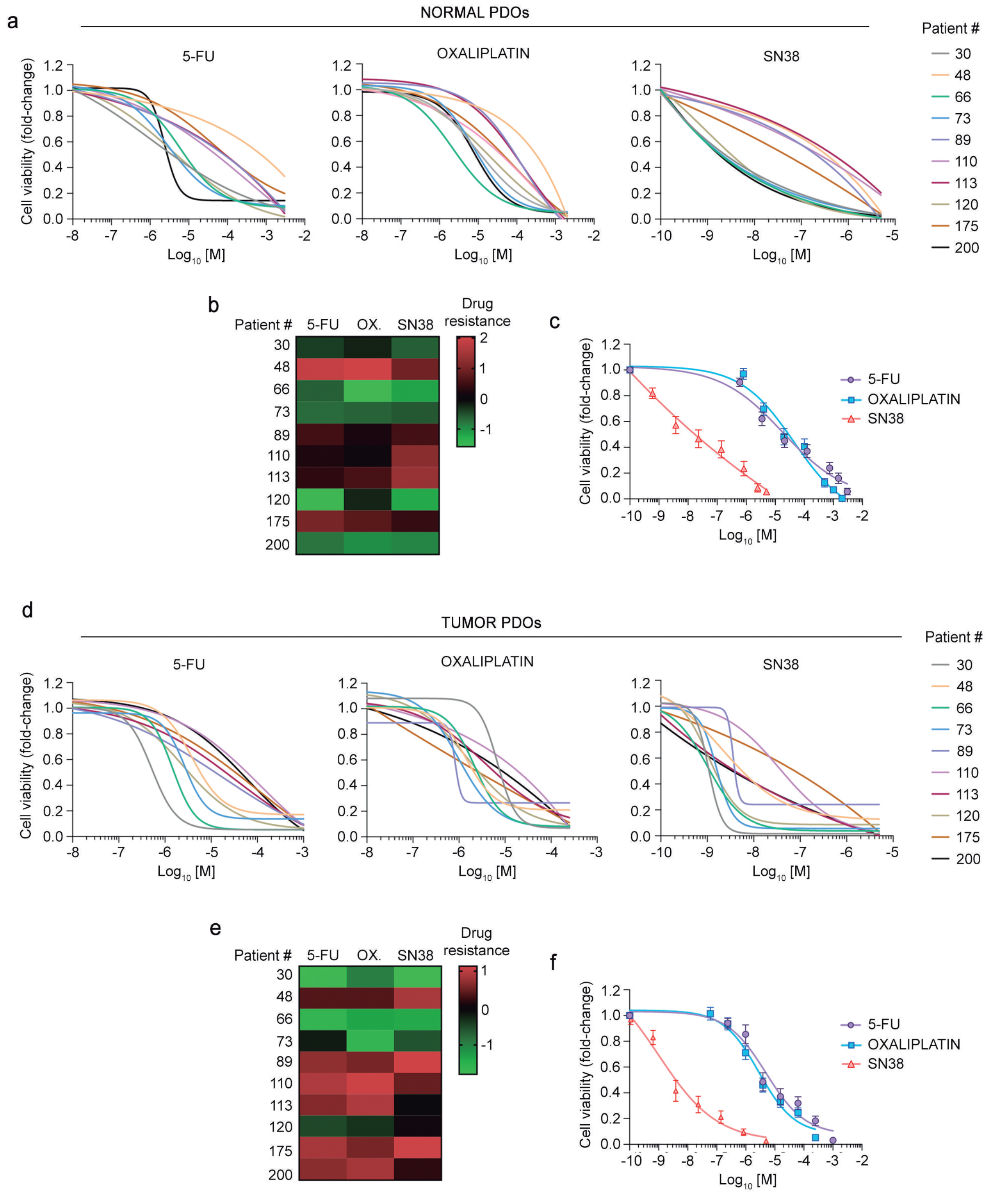


Supplementary Fig. 1: Response of colorectal normal and tumor PDOs to 5-FU, oxaliplatin and SN38. **a** Dose-response curves of 10 normal PDOs treated with 5-FU, oxaliplatin or SN38. Each line represents a patient. **b** Heatmap illustrating the z-score of drug resistance of normal PDOs from (**a**) for each drug treatment. **c** Dose-response curves (mean values +/- SEM) of normal PDOs used in (**a**). **d** Dose-response curves of 10 tumor PDOs treated with 5-FU, oxaliplatin or SN38. Each line represents a patient. **e** Heatmap illustrating the z-score of drug resistance of tumor PDOs from (**d**) for each drug treatment. **f** Dose-response curves (mean values +/- SEM) of tumor PDOs used in (**d**).
