## Supplementary Figure 2 for "Age-dependent chemotherapy response and vitamin D impact in paired patient-derived normal and tumor colorectal organoids"

Supplementary Fig. 2.

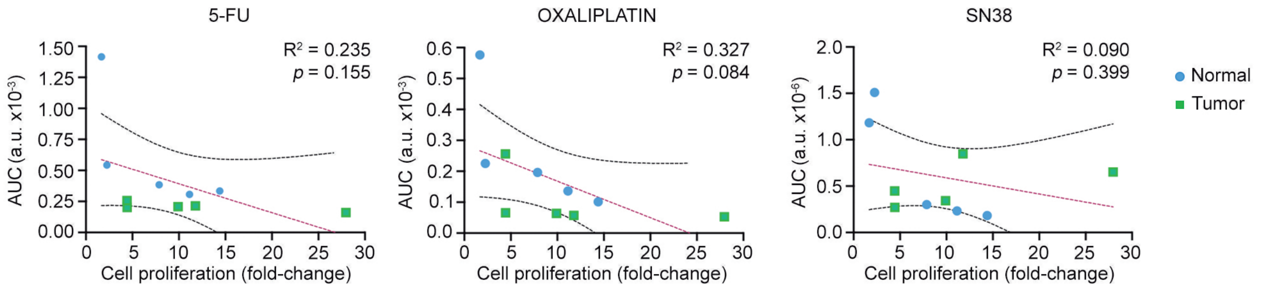


**Supplementary Fig. 2: Relationship between proliferation rates and drug cytotoxicity in colorectal normal and tumor PDOs.** Scattergrams showing the correlations between cell proliferation rates (assessed after 4 days in culture) and AUC values derived from the drug assays shown in Supplementary Fig. 1a and d. Each dot represents an individual normal (blue) or tumor (green) PDO treated with 5-FU, oxaliplatin or SN38. Correlation was evaluated by Pearson's test.
