## Supplementary Figure 3 for "Age-dependent chemotherapy response and vitamin D impact in paired patient-derived normal and tumor colorectal organoids"

Supplementary Fig. 3

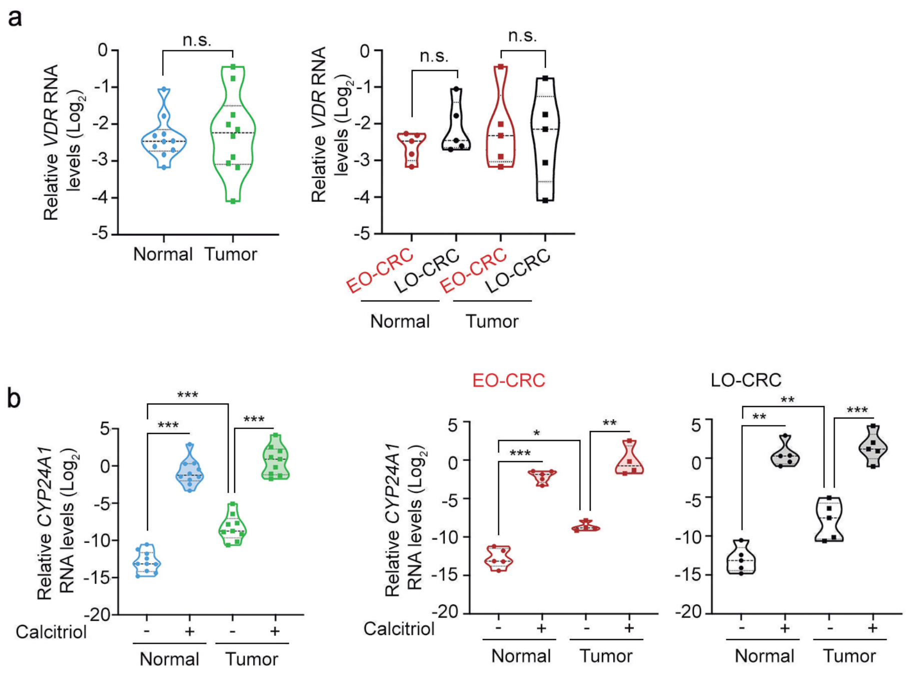


**Supplementary Fig. 3: Colorectal normal and tumor PDOs respond to calcitriol. a** Violin graphs of VDR RNA levels in normal and tumor PDOs. Overall (left panel) and age-grouped (right panel) PDOs. Statistical analysis was performed by Welch's t-test (n.s., non-significant). **b** Violin graphs of CYP24A1 RNA levels in normal and tumor PDOs treated for 96 h with 100 nM calcitriol or vehicle (ethanol). Overall (left panel) and age-grouped (middle and right panels) PDOs. One-way ANOVA with Sidak's post-test was applied for multiple comparisons (*P<0.05; **P<0.01; ***P<0.001).
