## Supplementary Figure 4 for "Age-dependent chemotherapy response and vitamin D impact in paired patient-derived normal and tumor colorectal organoids"

Supplementary Fig. 4.

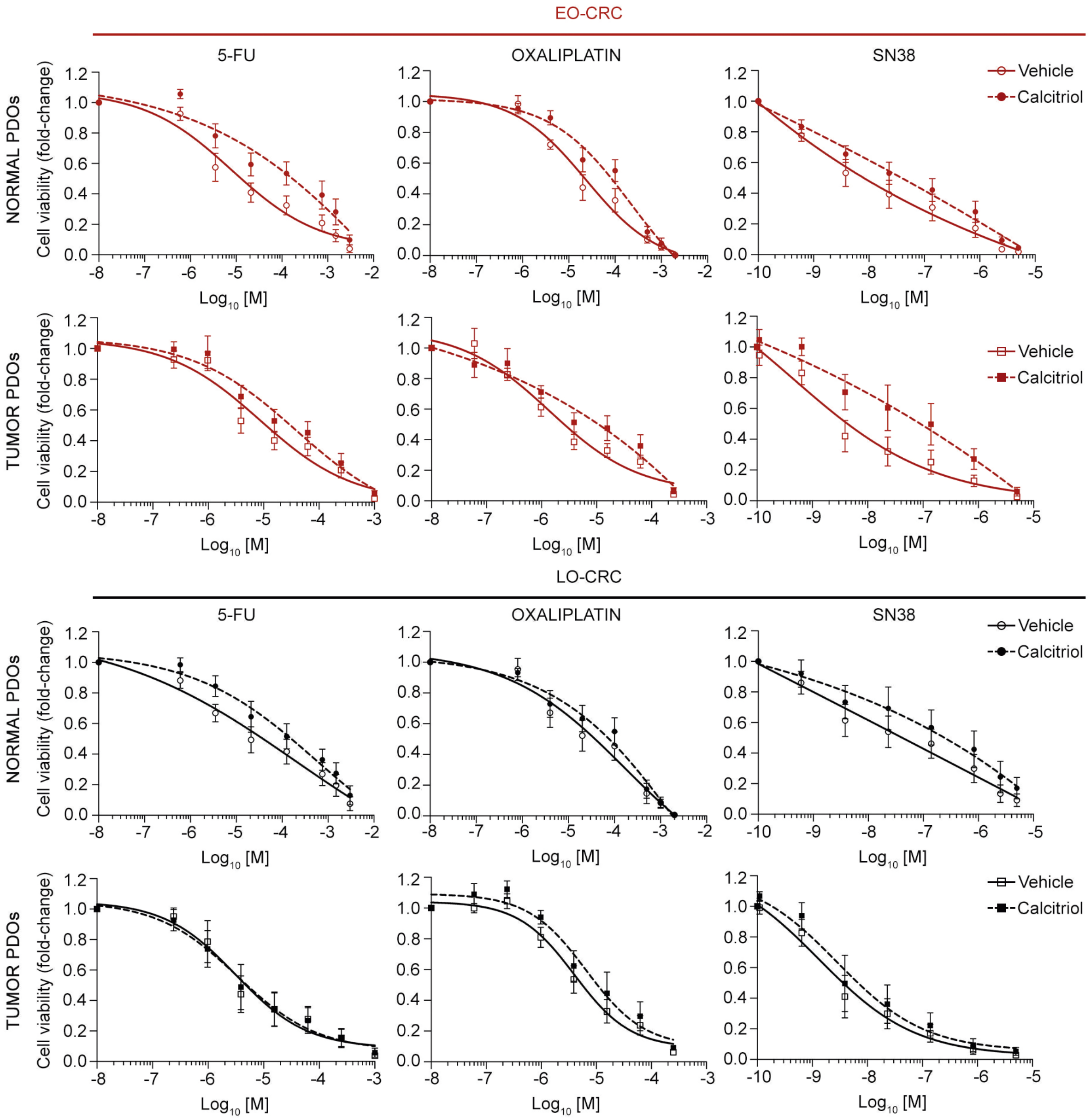


Supplementary Fig. 4: Effect of calcitriol on drug citotoxicity in EO-CRC and LO-CRC normal and tumor PDOs. Dose-response curves of normal and tumor organoids from EO-CRC and LO-CRC patients treated with 5-FU, oxaliplatin and SN38 in the presence of 100 nM calcitriol or vehicle (ethanol) for 96 h. Mean values +/- SEM are shown (n = 5).
