## Supplementary Table 1 for "Age-dependent chemotherapy response and vitamin D impact in paired patient-derived normal and tumor colorectal organoids"

**Supplementary Table 1: Clinicopathological characteristics of CRC patients whose biopsies were used to establish normal and tumor organoid cultures**

| Patient number | Sex | Age (years) | Classification | Tumor localization | T[a] | N[b] | M[c] | Drug assay | RNA-seq |
|---|---|---|---|---|---|---|---|---|---|
| 30 | Male | 81 | LO-CRC | Left colon | 4 | 1 | 0 | Yes | Yes |
| 48 | Male | 83 | LO-CRC | Left colon | 4 | 1 | 0 | Yes | No |
| 53 | Male | 81 | LO-CRC | Left colon | 3 | 0 | 0 | No | Yes |
| 57 | Male | 86 | LO-CRC | Left colon | 4 | 1 | 0 | No | Yes |
| 66 | Male | 76 | LO-CRC | Left colon | 3 | 0 | 0 | Yes | Yes |
| 73 | Female | 40 | EO-CRC | Left colon | 3 | 1 | 0 | Yes | Yes |
| 74 | Male | 84 | LO-CRC | Left colon | 3 | 0 | 0 | No | Yes |
| 89 | Male | 41 | EO-CRC | Right colon | 4 | 0 | 1 | Yes | Yes |
| 110 | Male | 57 | LO-CRC | Left colon | 3 | 0 | 0 | Yes | No |
| 113 | Male | 71 | LO-CRC | Left colon | 3 | 0 | 0 | Yes | No |
| 120 | Male | 45 | EO-CRC | Left colon | 3 | 0 | 0 | Yes | Yes |
| 122 | Male | 86 | LO-CRC | Left colon | 4 | 0 | 0 | No | Yes |
| 175 | Female | 46 | EO-CRC | Rectum | 4 | 2 | 0 | Yes | No |
| 183 | Male | 49 | EO-CRC | Left colon | 1 | 1 | 0 | No | Yes |
| 200 | Male | 48 | EO-CRC | Left colon | 4 | 1 | 0 | Yes | Yes |

**T[a]**: Size or direct extent of the primary tumor; **N[b]**: degree of spread to regional lymph nodes; **M[c]**: presence of distant metastasis (www.uicc.org/resources/tnm).
