## Supplementary Table 2 for "Age-dependent chemotherapy response and vitamin D impact in paired patient-derived normal and tumor colorectal organoids"

Supplementary Table 2: Composition of culture media for colorectal normal and tumor PDOs.

| **Reagent** | **Concentration** | **Source** | **Normal medium** | **Tumor medium** |
|---|---|---|---|---|
| Advanced DMEM/F12 | | Thermo Fisher Scientific | 50% | 100% |
| HEPES | 10 mM | Thermo Fisher Scientific | + | + |
| Glutamax | 10 mM | Thermo Fisher Scientific | + | + |
| B27 supplement | 1X | Thermo Fisher Scientific | + | + |
| N2 supplement | 1X | Thermo Fisher Scientific | + | + |
| Nicotinamide | 10 mM | Sigma-Aldrich | + | - |
| N-acetyl-L-cysteine | 1 mM | Sigma-Aldrich | + | + |
| Primocin | 0.1 µg/mL | Invivogen | + | + |
| L-WNR (Wnt3A, Noggin, RSPO3) conditioned medium | | Custom made (Ref. #44) | 50% | - |
| RSPO1 | 1 µg/mL | Sinobiological | + | - |
| $PGE_2$ | 0.02 µM | Sigma-Aldrich | + | + |
| EGF | 50 ng/mL | Peprotech | + | + |
| Gastrin | 1 µg/mL | Tocris | + | + |
| Noggin | 0.1 µg/mL | Sinobiological | + | + |
| Y-27632 | 10 µM | Tocris | + | + |
| LY2157299 | 1 µM | Axon-Medchem | + | + |
| SB202190 | 10 µM | Sigma-Aldrich | + | + |
